# Genomic insights into red squirrels in Scotland reveals loss of heterozygosity associated with extreme founder effects

**DOI:** 10.1101/2024.07.10.602565

**Authors:** Melissa M. Marr, Emily Humble, Peter W. W. Lurz, Liam A. Wilson, Elspeth Milne, Katie M. Beckmann, Jeffrey Schoenebeck, Uva-Yu-Yan Fung, Andrew C. Kitchener, Kenny Kortland, Colin Edwards, Rob Ogden

## Abstract

Remnant populations of endangered species often have complex demographic histories associated with human impact. This can present challenges for conservation as the genetic status of these populations are often a-typical of natural populations, and may require bespoke management. The Eurasian red squirrel, *Sciurus vulgari*s (L., 1758), is endangered in the UK. Scotland represents a key stronghold, but Scottish populations have been subjected to intense anthropogenic influence, including wide-spread extirpations, reintroductions and competition from an invasive species. This study examined the genetic legacy of these events through whole genome resequencing of 106 red squirrels. Using SNP and genotype likelihood datasets, previously undetected population structure and patterns of gene-flow were uncovered. One off-shore island, three mainland Scottish populations, and a key east-coast migration corridor were observed. An abrupt historical population bottleneck related to extreme founder effects has led to a severe and prolonged depression in genome-wide heterozygosity, which is amongst the lowest reported for any species. Current designated red squirrel conservation stronghold locations do not encompass all existing diversity. These findings highlight the genetic legacies of past anthropogenic influence on long-term diversity in endangered taxa. Continuing management interventions and regular genetic monitoring are recommended to safeguard and improve future diversity.

## 1. Introduction

Contemporary populations of endangered mammals often have complex demographic histories that present challenges for conservation management. The last 500 years has seen unprecedented and accelerating biodiversity loss (Barnosky, 2008; Dirzo et al., 2014) coupled with widespread destruction and fragmentation of habitats (Haddad et al., 2015; Maxwell et al., 2016; Püttker et al., 2020). This has reduced many populations to remnant, disjunct, habitat patches, placing them at greater risk of both local and global extinction (Crooks et al., 2017; Haddad et al., 2015; Hanski, 2015). This situation is further exacerbated in some regions by the accidental or deliberate release of species that become invasive, one of the leading drivers of extinctions via resource competition and introduction of novel diseases (Clavero and Garciaberthou, 2005; Crowl et al., 2008; Gallien and Carboni, 2017).

Human-mediated movement of fauna has undoubtedly occurred for millennia (Hofman and Rick, 2018), while reintroductions and other forms of conservation translocation are now commonplace tools in the preservation and genetic of threatened taxa (Frankham, 1996; Seddon et al., 2007; Taylor et al., 2017). As a result, populations of some endangered species have experienced inter-and intra-specific admixture in addition to other long-term demographic changes, which may include population declines and/or fluctuations, local extirpations and increased isolation (*e.g.,* Florida panther *Puma concolor coryi* (Johnson et al., 2010), red wolf *Canis rufus* (Sacks et al., 2021) and alpine marmots *Marmota* spp. (Kerhoulas et al., 2015)).

Populations with such complex demographic pasts may represent artificially admixed assemblages and exhibit atypical genetic histories compared to large, unmixed, natural populations. They are often associated with poor genetic health, exhibiting high levels of inbreeding and low genetic diversity due to past population bottlenecks and small numbers of founding individuals (Frankham et al., 2010; Keller, 2002). This can lead to genetic erosion, loss of adaptive potential and increased extinction risk (Leroy et al., 2018; Mathur and DeWoody, 2021; Willi et al., 2006). Despite this, such populations can have huge conservation importance as they often represent the sole remnants of that species in a particular region (*e.g.* Iberian lynx *Lynx pardinus* (IUCN, 2014), Ethiopian wolf *Canis simensis* (Mooney et al., 2023) and, Pacific pocket mouse *Perognathus longimembris pacificus* (Wilder et al., 2022)). As biodiversity and habitat losses continue apace, it is essential that fragmented and genetically depauperate populations are managed effectively in the face of increased threats to their long-term survival.

In Britain, the Eurasian red squirrel (*Sciurus vulgaris* L., 1758) exemplifies a species with a complex demographic history related to changes in land use and cultural attitudes (Harvie-Brown, 1881a, 1881b, 1880; Holmes, 2015). Furthermore, it has become a text-book icon of conservation science due to its 20^th^ century (and continuing) replacement by the introduced North American grey squirrel *Sciurus carolinensis* (G., 1788 (Gurnell et al., 2004; Gurnell and Pepper, 1993; Rushton et al., 2006)), which has arguably become one of the best-known examples of a native species being supplanted by an invasive competitor (Gurnell et al., 2004; Wauters et al., 2023). While habitat loss and fragmentation contribute to current population declines, the major driver of contemporary replacement is ecological and disease-mediated competition with the introduced grey squirrel, which can asymptomatically carry a virus novel to red squirrels, squirrelpox virus SQPV (Rushton et al., 2006; Sainsbury et al., 2000; Tompkins et al., 2003).

Red squirrels are now almost entirely absent from mainland England and Wales. Scotland represents a stronghold for the species in Britain, harbouring over 80% of the remnant population (*c*. 239,000 individuals (Mathews et al., 2018)). However, historical Scottish populations experienced dramatic range contractions and local extirpations prior to the introduction of *S. carolinensis* (Harvie-Brown, 1881a, 1881b, 1880). Some authors suggest the occurrence of a near-extinction event around the 18th century linked to low forest cover (Harvie-Brown, 1881a, 1881b; Kitchener, 1998; Ritchie, 1920). This was followed by a period of unofficial and unmanaged restocking across Scotland via introductions from populations in Central/NW Europe and England from the late 1700s (Harvie-Brown, 1881a, 1880, 1880; Ritchie, 1920). Numbers of introduced individuals are unknown, but this likely involved introduction of small numbers of animals from disparate origins, into several Scottish locations over a number of decades. It is probable that modern populations represent admixed composites of the remnant Scottish population with English and European additions.

Red squirrels are a conservation priority in Scotland with protection under the Nature Conservation (Scotland) Act 2004 and are listed in the Scottish Biodiversity List (https://www.nature.scot/doc/scottish-biodiversity-list). A suite of conservation measures are currently in place for red squirrels in Scotland under the Species Action Framework, including forestry management and grey squirrel control (Gaywood, 2016). Chief among these is the creation by the Forestry Commission (now Scottish Forestry and Forestry and Land Scotland) of 19 ‘stronghold’ woodlands, where management is focused on creating an ecological advantage for red squirrels (Forestry Commission Scotland, 2012). Translocations of red squirrels are also taking place *ad hoc* in the north of the country, with red squirrels from Moray and Inverness being moved to areas in the north-west Highlands to restock this area after a long interval of absence (Dennis, 2012). Movement of red squirrels to re-establish extirpated populations in Britain has been a widely used approach in red squirrel conservation, with notable successes (*e.g.,* Anglesey, Wales (Shuttleworth and Halliwell, 2016)) and failures (Sainsbury et al., 2020). Disease, particularly squirrelpox, has been confidently linked to translocation failure (Sainsbury et al., 2020), but the role of genetic factors has never been fully explored due to the lack of informative data.

Despite the importance of Scottish populations to the survival of red squirrels in Britain, little is known about their genetic status and population structure. Analysis of English and Welsh populations has uncovered patterns of low within-population diversity and high among-population differentiation, but with relatively little phylogeographical structure, indicative of serial translocations, severe historical population bottlenecks and little contemporary gene flow (Barratt et al., 1999; Hale et al., 2004; Ogden et al., 2006). Under a scenario where effective population size is small and immigration rates low, deleterious and recessive alleles will tend to accumulate as genome-wide homozygosity (including loci affecting fitness) decreases under strong drift and weak selection (Frankham, 1996; Frankham et al., 2010; Keller, 2002). Such scenarios can also be precipitated by small numbers of founding individuals*, i.e.* founder effects (Szucs et al., 2017), and confounded when dispersal is limited, either by natural or artificial landscape barriers or when populations are supressed by competitors. Long-term connectivity is key for the maintenance of gene flow and avoidance of genetic drift between fragmented populations (Frankham, 1996; Lowe and Allendorf, 2010).

Low diversity, high inbreeding and population fragmentation could reduce the ability of Scottish red squirrel populations to cope with stochastic events and disease outbreaks. Moreover, it could impede the expansion of existing Scottish populations and reduce their utility to act as founders for new populations, thereby jeopardising red squirrel recovery in Britain.

To address these key conservation concerns, our study undertook the first genome-wide assessment of red squirrels across Scotland in order to address the specific questions, *i*) how are populations genetically structured? and, *ii*) what are the levels of inbreeding and genome-wide diversity, and how are these partitioned among populations and landscapes? Our results are discussed both within the context of applied red squirrel-specific management and their wider applicability to the conservation of endangered species facing these common threats.

## 2. Materials and Methods

### 2.1. Sample collection

Specimens were sourced from frozen red squirrel carcasses that had been collected as part of a long-running disease surveillance program at the Royal (Dick) School of Veterinary Science, University of Edinburgh as well as from National Museums Scotland (NMS). The opportunity arose to include samples from a remnant English population in Formby (Sefton). Scottish and English populations may have different demographic histories, but very few English mainland populations remain (Mathews et al., 2018). Therefore, the inclusion of samples from Formby provided a means to include a comparative English outgroup (Fig.1., Table S1).

**Figure 1.**
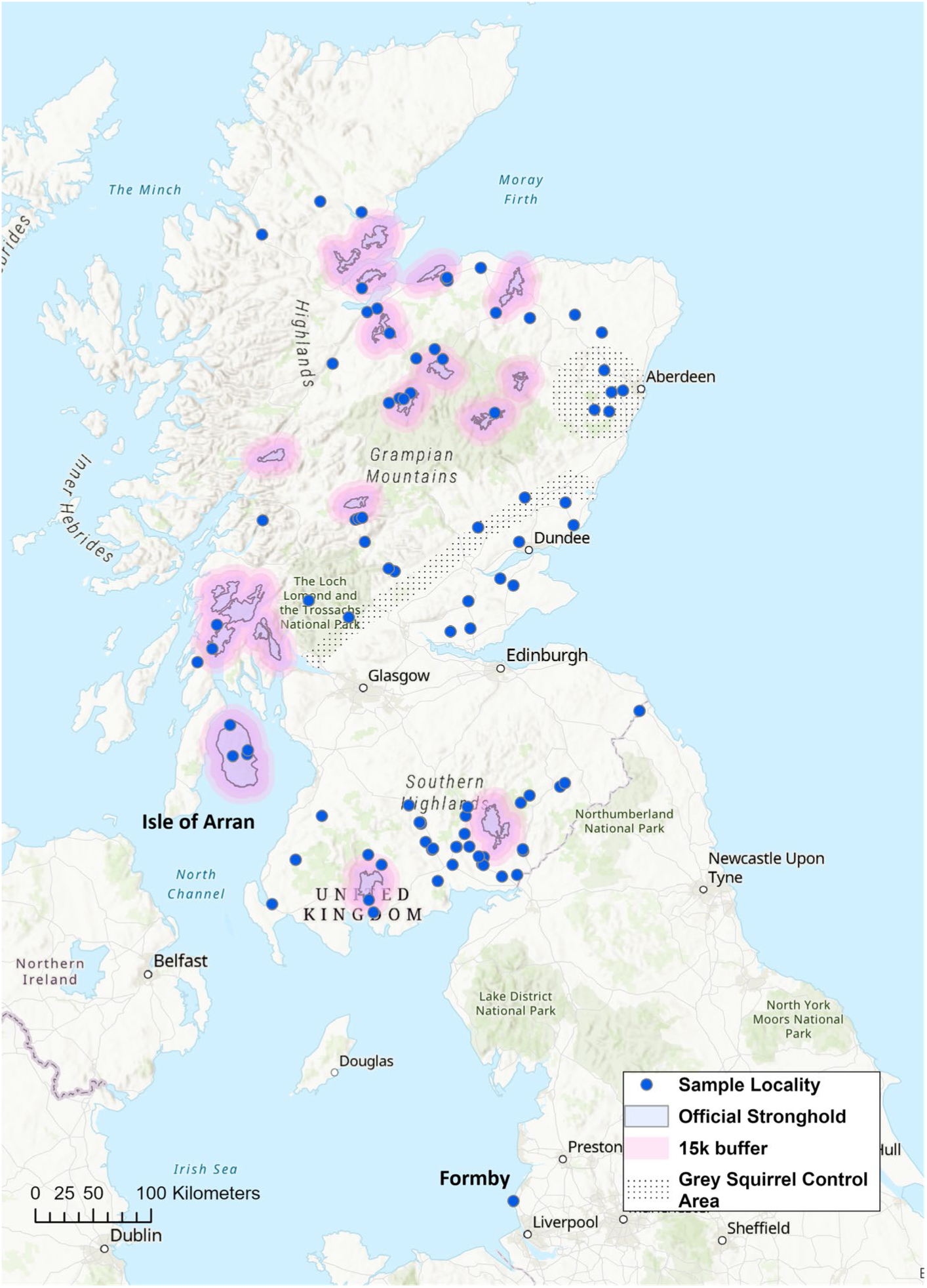
Sample distribution of red squirrels included in the study. Sample localities are the locations where dead red squirrels were found. Official stronghold areas are shown in pink with 15km buffer inclusion zones. Note the absence of specimens from the Central Belt, where there is a discontinuity in red squirrel distribution separating populations in the south of Scotland from those in the north. Most strongholds are located on mainland Scotland, aside from one on the Isle of Arran.

Tissues were sampled from whole organs (kidney, liver, heart) while museum samples were from uterine and genital tissues (Table S1). All tissue sampling was performed either from thawed carcases (R(D)SVS)) or thawed tissues (NMS), with all surfaces and instruments cleaned with 10% bleach solution between individuals to avoid contamination. Data were filtered to select squirrels that had at least a six-digit OS grid reference. Specimens were added as point data to a UK basemap using ESRI ArcGIS Pro v2.8, with layers added to show official red squirrel strongholds and grey squirrel cull/buffer zones (Fig. 1). The final database consisted of 94 specimens from Scotland and 12 individuals from Formby, giving a total of 106 individuals. (Fig. 1, Table 1). Specimens were grouped into six putative populations based on geographical location for *a priori* hypothesis testing as follows: HIG - Highlands and Moray, NE - north-east Scotland inc. Aberdeen City and Shire, CEN – central Scotland inc. Fife, Perthshire and Argyll, ARR – Isle of Arran, SW Scotland – inc. Dumfries and Galloway and south Strathclyde, BOR – Scottish Borders, FOR - Formby (Fig. 1, Table 1). These areas have unique and disparate population histories, with varying numbers of founders from a variety of (often undocumented) origins, as well as variations in habitat availability and presence of grey squirrels. To compare diversity inside and outside stronghold areas, squirrels were also subdivided by stronghold grouping.

**Table 1.**
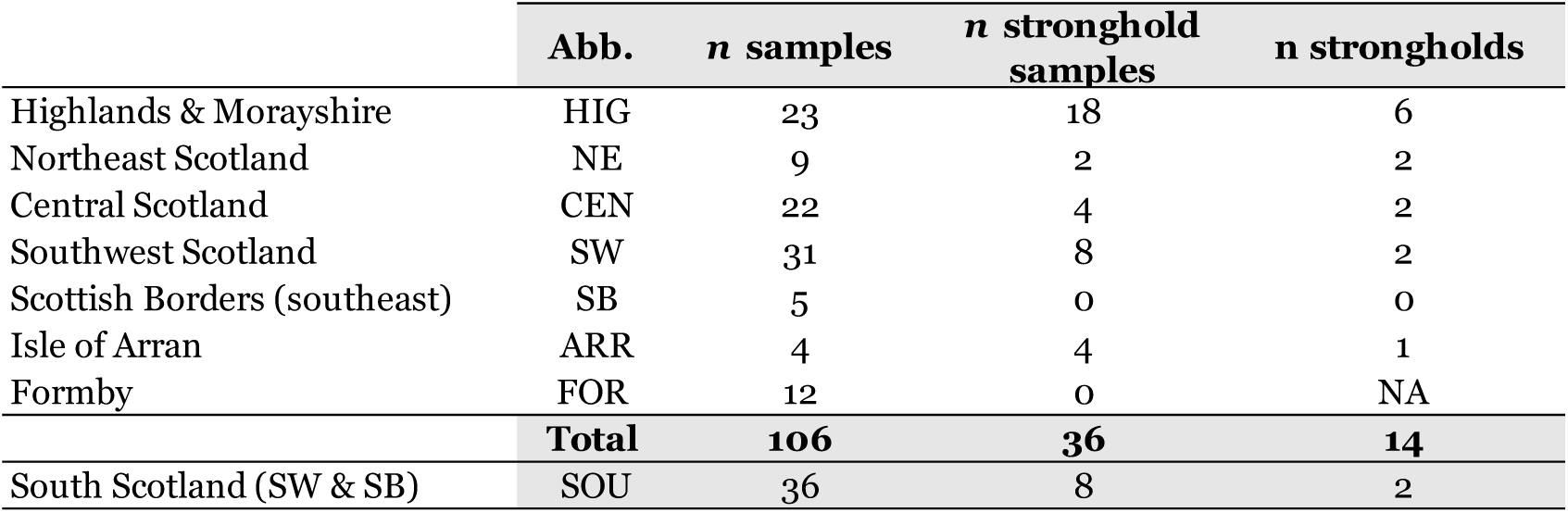
Sample numbers and stronghold distribution. Fourteen of the nineteen designated red squirrel Scottish stronghold areas are represented in the study, with a clear geographical bias. Most stronghold areas are located in the Highlands and NW regions. The Isle of Arran represents one stronghold. The English population Formby is not part of the Scottish stronghold initiative.

|  | Abb. | n samples | n stronghold samples | n strongholds |
| --- | --- | --- | --- | --- |
| Highlands & Morayshire | HIG | 23 | 18 | 6 |
| Northeast Scotland | NE | 9 | 2 | 2 |
| Central Scotland | CEN | 22 | 4 | 2 |
| Southwest Scotland | SW | 31 | 8 | 2 |
| Scottish Borders (southeast) | SB | 5 | 0 | 0 |
| Isle of Arran | ARR | 4 | 4 | 1 |
| Formby | FOR | 12 | 0 | NA |
|  | **Total** | **106** | **36** | **14** |
| South Scotland (SW & SB) | SOU | 36 | 8 | 2 |

### 2.2. DNA Extraction and Sequencing

DNA was extracted from c. 50 mg of tissue using the automated Promega Maxwell® RSC instrument in conjunction with the Maxwell® RSC PureFood and GMO Authentication Kit. Double-stranded, single-index, libraries were constructed by Azenta using the NEB Ultra II DNA Library Preparation Kit. Paired-end (PE) 150bp sequencing was then conducted on an Illumina Novaseq for a desired c. 5X sequence coverage per sample.

### 2.3. Whole Genome Bioinformatic Processing

#### 2.3.1. Read alignment

The *S. vulgaris* reference assembly (Mead et al., 2020) version mSciVul1.2, GCA_902686455.2, was downloaded from the NCBI, before being modified to include only the nineteen autosomes. Reference genome indexing was performed with the Burrows-Wheeler aligner (BWA v.2.1.0 (Li and Durbin, 2009)). Raw, paired-end, Illumina reads were quality and adapter trimmed with Trim Galore v.0.6.4 (https://github.com/FelixKrueger/TrimGalore) and reads were aligned to the reference genome using the BWA-MEM algorithm within BWA-Kit (Li and Durbin, 2009), which additionally marks duplicates and adds read-group information. Alignments were sorted and filtered with SAMtools v.1.10 (Danecek et al., 2021) to remove unmapped reads (flag *–F 4*) and reads with mapping quality < Q30. PCR duplicates were removed with Picard – MarkDuplicates (http://broadinstitute.github.io/picard).

#### 2.3.2. Calling SNPs and genotype likelihoods

Read coverage distribution was quantified in ANGSD v.0.941-6-g67b6b3b-dirty (Korneliussen et al., 2014), using the *–doDepth* function with *–mapMapQ* 30*, -doQdists* 1*, -doCounts* 1 and *–maxDepth* set to 800. The ngsTools (Fumagalli et al., 2014) R script, plotQC.R, was used visualise depth distributions and calculate the upper and lower 5^th^ percentiles, which were removed from SNP and genotype likelihood (GL) calculations. SNP calling and GL estimation runs were performed in ANGSD, using the GATK (Poplin et al., 2017) model for genotype calling (*-GL 2*) and the beagle likelihood model (*-doGLF* 2). The reference allele was set as the major allele (*-doMajorMinor* 4) and minimum mapping quality was set to 30 (*-minMapQ* 30), minimum base score quality to 30 (*-minQ* 30) and the p-value cut-off for SNPS set to p < 1 * 10^-6^. For analyses that required direct inference from high quality SNPs, a second ANGSD run was undertaken with the minimum coverage per site required for a call increased to 4X and only sites with no missing data were considered.

#### 2.3.3. Estimating the folded site-frequency spectrum (SFS)

Folded site frequency spectra (SFS, *-fold* 1) were estimated for each population and for all pairs of populations (joint, 2D, SFS), using winsfs (Rasmussen et al., 2022), based on the per-population site allele frequencies (SAFs, *-doSAF* 1). The reference genome was specified as the ancestral genome.

### 2.4. Data Analysis

#### 2.4.1. Population structure

Genotype likelihoods were pruned for sites in linkage disequilibrium (LD), to ensure that population structure was not driven by highly correlated loci. Data were first down-sampled to 1 in every 20 SNPs for computational efficiency. The ngsLD v.1.1.0 package (Fox et al., 2019) was used to calculate pairwise measures of LD, assuming a max distance of 1Mb between SNPs, before pruning with the prune_graph.pl script (max distance 5kb, minimum weight 0.5). Pruned, unlinked, GLs were used to generate a covariance matrix with PCAngsd (Meisner and Albrechtsen, 2018), with a minor allele frequency filter (MAF) of 0.05. The same dataset was then used to examine admixture proportions per individual using NGSadmix (Skotte et al., 2013). K was set to range between 1 and 20, with 10 replicates per value of K. Assessment of the plateau of the likelihood curve and the deltaK method (Evanno et al., 2005) were used to determine the most likely value of K. PCA and admixture outputs were plotted using custom scripts in R v.4.2.2 (R Core Team, 2021).

#### 2.4.2. Fast estimation of effective migration surfaces (FEEMS)

Spatial analysis of gene flow was explored using FEEMS v. 1.0. (Marcus et al., 2021), using the SNP dataset with 4x average minimum coverage per site and no missing data. Plink v.1.90 was used to apply a MAF filter of 0.05 and to prune SNPs in high LD (win 50, step 10, threshold 0.1). The resulting matrix of unlinked, biallelic SNPs was used as FEEMS input. Populations were then assigned to vertices of a 5km triangular discrete global grid (DGG) before estimation of effective migration via a penalised-likelihood λ framework. Values of λ=2, 5 and 10 were plotted to explore the effect of lambda on structure; in general, lower values tend to lead to stronger population structure and overfitting of the model.

#### 2.4.3. Population divergence – fixation index Fst

Population differentiation was investigated by estimating the pairwise fixation index, F_ST_, for all population pairs, where populations were defined on the basis of the PCA, admixture and migration results. Per-site pairwise estimates of weighted F_ST_ were then calculated by the realSFS program using the folded 2D SFS. Weighted per-site F_ST_ estimates were calculated using the Hudson estimation (Bhatia et al., 2013), specified using the *-whichFST* 1 option (recommended for low coverage data), and the sliding-window approach with window sizes of 50-kb and 10-kb intervals. Global weighted FST was calculated using the realSFS F_ST_ function for all population pairs.

#### 2.4.4. Neutrality and Diversity

Thetas and Tajima’s D neutrality statistics were calculated from the population-level folded SFS using a sliding window approach. The thetaStat function was used with the *–do_stat* option with a window size of 50kbp (*-win* 5000) and a step size of 10kbp *(-step* 10000). Final values for both statistics were calculated by dividing the output by the effective number of sites in the. pestPG file. Estimates of individual genome-wide heterozygosity were calculated from the individual SFS using the *-realSFS* function.

#### 2.4.5. Runs of homozygosity (ROH)

Inbreeding was investigated by calculating runs of homozygosity (ROH) in Plink v.1.90 (Chang et al., 2015) using the min 4X coverage dataset with no missing data using the Plink --homozyg function. A minimum length of 1Mb (*--homozyg-kb* 1000), minimum of 50 SNPs *(--homozyg-window-snp* 50) and only 1 heterozygous site *--homozyg-het* 1 were required to call a ROH. Additional parameters were: *--homozyg-window-missing* 3, *--homozyg-window-threshold* 0.05, *--homozyg-gap* 1000, *--homozyg-density* 70. A minor allele frequency filter of 0.05 was also added (*--maf* 0.05). Individual inbreeding coefficients (F_ROH_) were calculated as the sum of ROH lengths divided by the length of the autosomal genome (2.53Gb), with bin lengths of 1-2.5Mb, 2.5-5Mb, 5-7.5Mb and > 7.5Mb.

#### 2.4.6 Recent demographic history, N_e_

Changes in recent demography were estimated for each population, using changes in effective population size, *N_e,_* up to 200 generations. *N_e_* estimations were performed with GONE (Santiago et al., 2020), based on patterns of linkage disequilibrium (LD). This was directly estimated from SNPs, using the SNP dataset (section 2.5.2) with a MAF filter of 0.05. Data were converted to Plink format and the programme was run 5 times per population, with the average values plotted using a custom R script. To convert generations to calendar years, we estimated the average squirrel generation time as three years. This is based on life history information (Lurz et al., 2005), interpreted from the ecology of the species.

#### 2.4.7. Stronghold diversity

To compare genetic diversity (measured by individual heterozygosity) within and outside of stronghold areas, data were partitioned by population and then grouped as within or outside a stronghold area (Table 1). There are eighteen official mainland strongholds and one offshore-island stronghold on the Isle of Arran (Fig. 1, (Slade et al., 2021)). Individuals within the designated borders of the strongholds are under-represented in the dataset due to geographical remoteness. Therefore, we decided to include 15km buffer zones around strongholds to increase the inclusion criteria and, consequently, the sample size, as contiguous/neighbouring areas likely benefit from the stronghold management effect (Fig. 1). This effectively led to the merger of neighbouring strongholds in some areas. A series of two-tailed, independent, t-tests were then performed to test for significant differences in heterozygosity between groups that had adequate sample sizes. This was repeated for pooled population data.

#### 2.4.8. Mitochondrial genome assembly and analysis

Trimmed reads were mapped against the *S. vulgaris* reference mitogenome (accession LR822068.1), using the same methods and post processing as for the nuclear genome. Consensus sequences were constructed using MUSCLE (Edgar, 2004) in the Geneious Platform v.8 (Kearse et al., 2012), with a minimum of 5 reads coverage per site for a call, before extracting and concatenating the two rRNAs, 22 tRNAs, 13 protein-coding genes and the D-Loop. A median-joining (MJ) phylogenetic network (Bandelt et al., 1999) was generated in PopART (Leigh and Bryant, 2015). Estimation of mitochondrial diversity was estimated via haplotype (Hd) and nucleotide diversity (π) in DNAsp v6. (Rozas et al., 2017).

## 3. Results

### 3.1. Illumina short-read sequencing and read alignment

After read alignment to the nineteen autosomes of the *Sciurus vulgaris* reference genome (GCA_902686455.2 (Mead et al., 2020)), mean coverage ranged from 3.7X to 7.5X (SI_Sample_Database). Two samples, R232.98 (NMS.Z.2000.195.46) and R235.98 (NMS.Z.2000.195.49), generated unexpectedly large amounts of data and had 10X and 22.5X coverage, respectively. Mean x-fold coverage after read alignment to the reference mitogenome resulted in a minimum of 60X coverage. The low coverage ANGSD run produced 9,324,158 SNPs, while the second run, excluding sites < 4X with no missing data produced 123,052 SNPs (stored in Harvard Dataverse repository https://doi.org/10.7910/DVN/CK1ILL). A density plot of the 9,324,158 SNPs showed a largely uniform distribution, but with some minor high-density areas and two notable SNP hotspots on chromosomes 13 and 16 (Fig. S1). The effect of removing these areas on the site frequency spectrum was explored, but found to be minimal (Fig. S2).

### 3.2. Mitochondrial genome

Consensus sequences of the mitogenome were between 16,511 to 16,516bp in length, consistent with published sequences for this species. The final trimmed alignment was 16,454 bp with 145 polymorphic (informative) sites. Haplotype diversity was moderate (*Hd*: 0.89), but nucleotide diversity was low (*π* = 0.00016). Out of 106 sequences, 23 haplotypes were detected (svm01-svm23, Fig. 2, Table S1). The most common haplotype was svm10 (n = 25), which was associated almost exclusively with the Highlands and NE Scotland region (one occurrence in the Central population), followed by svm06 (n = 14) found predominantly (but not solely) in the central region, and svm09 (n = 14) found only in areas south of the Central Belt (Fig. 2, Table S1).

**Figure 2.**
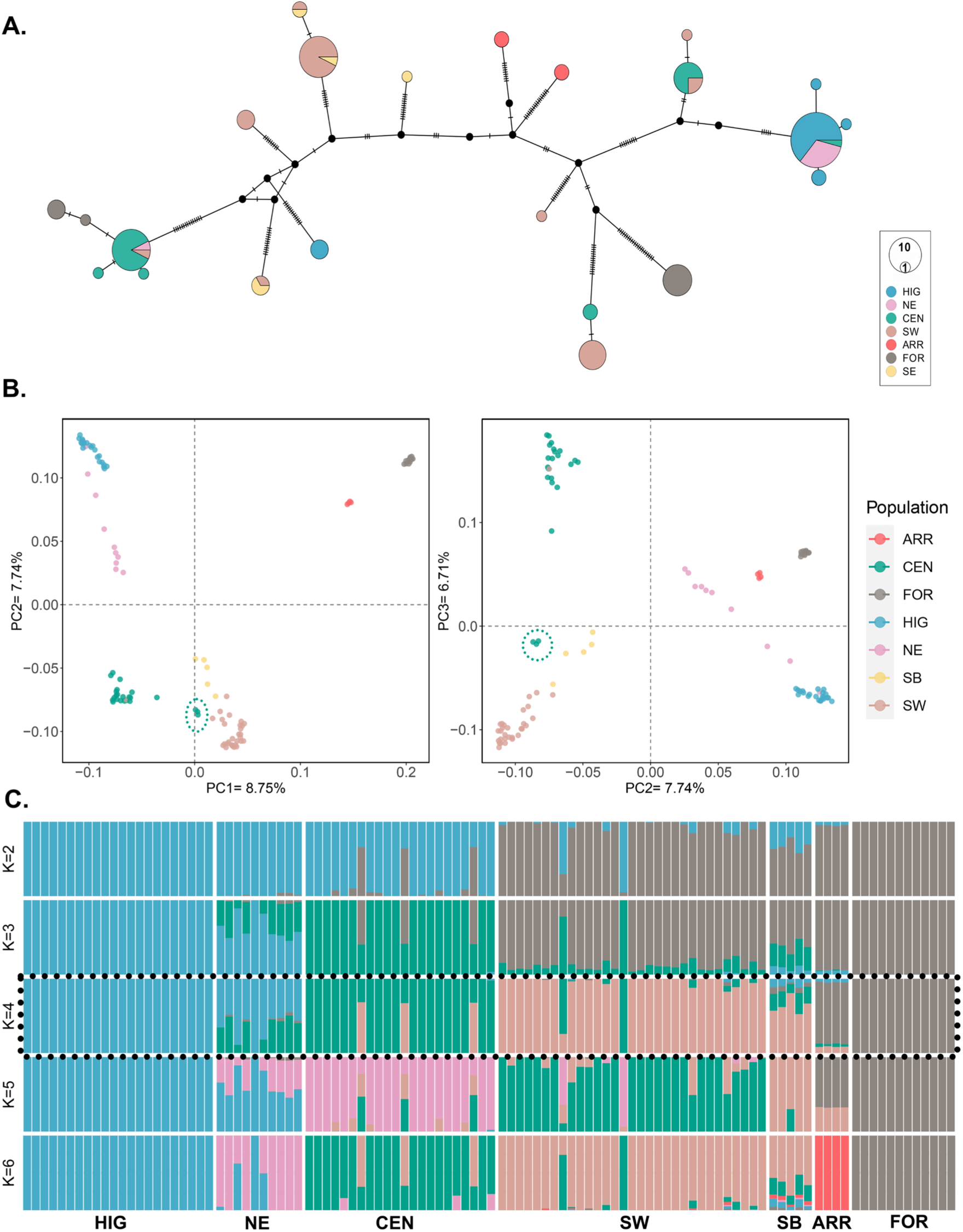
Population Structure. **A:** Mitochondrial genome haplotype network: Medium-joining (MJ) network for Scottish and Formby red squirrels. Branch lengths are drawn to scale and nodes are proportional to haplotype frequencies. 23 haplotypes were present in the dataset of 106 red squirrels with *Hd* = 0.89, *μ* = 0.00016. **B:** Principal components analysis (PCA). **C:** Admixture analysis. Admixture and PCA were performed on 775,219 genotype likelihoods (GLs). Both analyses suggest broad population divisions into three mainland populations (HIG + NE, CEN, SW + SB), and one population composed of ARR + FOR. Arran and Formby do not constitute a single population due to geographical separation, and the NE is considered separate in this study due to high levels of admixture. Green dashed circles in the PCA indicate the Argyll and Bute individuals that are geographically within the Central population but which appear genetically more similar to the SW squirrels.

Despite the geographical trends in haplotype frequencies, the overall pattern showed a distinct lack of phylogeographical structure, with highly divergent haplotypes observed at all localities. Eight haplotypes were unique and nine were present only in low frequencies (≥2≤5), with these low-frequency haplotypes distributed throughout the study region and associated mainly with the Scottish mainland populations (Fig. 2.). Most regions shared haplotypes with other areas, except the Isle of Arran. This population had two haplotypes, svm12 and svm15, which were unique to the island, but were highly divergent from each other, with 27 mutations between them. Formby, an English mainland ‘island’, had three haplotypes, which were not shared by any of the Scottish populations. The most common of which was smv13 (n=8). This haplotype was highly divergent from the other Formby haplotypes svm07 (n=3), and svm19 (n=1)), that were most similar to those found in the Central region (Fig. 2.)

### 3.3. Autosomal population structure, admixture and geneflow

In contrast to mitochondrial DNA, autosomal genotype likelihoods (GLs) showed clear patterns of population subdivision, admixture and migration that were highly consistent across analyses (Fig. 2). After MAF filtering (0.05) and pruning for sites in high LD, the PCA and admixture dataset consisted of 775,219 GLs. The PCA showed population groupings concordant with geographical structure (Fig. 2, Fig. S3). Principal component 1 (8.75%) separates Arran and Formby from the mainland Scottish populations. Along PC2 (7.74%), three geographical groups are apparent in the mainland Scottish populations: a northern group consisting of the Highlands and NE Scotland (with internal substructure), a Central Scotland group and a South Scotland group, consisting of the SW plus the Scottish Borders (also with internal substructure). These patterns remained after subsampling groups for equal sample sizes (Fig. S4).

The likelihood curve and delkaK unambiguously indicated K=4 as the most likely number of populations in the NGSadmix analyses (Fig. S5). These corresponded with the PCA groupings, suggesting a Highlands + NE group, a Central Scotland group, a Southern group (SW + Borders) and a population consisting of the Isle of Arran + Formby. The majority of individuals in the NE group were admixed between the northern Highlands group and the Central group to the south, suggesting that the NE is a contact zone. Three individuals from Argyll and Bute showed a genetic profile more typical of the southern population despite being geographically located within the Central group (Figs. 1,2). At higher values of K, the Scottish Borders (south-east) population appear separate from the SW population, and at K=6, ARR and FOR are distinguished.

After pruning the higher quality SNP dataset (min 4X, no missing data) for LD, the remaining number of SNPs for FEEMS analysis was 10,962. Spatial visualisation of red squirrel migration highlighted the lack of gene flow between populations north and south of the Scottish urban Central Belt (Fig. 3, Fig. S6). North-east Scotland, and the east coast, emerges as a key area for red squirrel gene flow between the Central and Highland populations. The three individuals, which fit the genetic profile of the southern population while being located in the Central region, are shown here as an isolated group on a peninsula in Argyll and Bute. There is a complete lack of migration along the west coast and indications that geneflow is somewhat reduced between the south-west and south-east regions.

**Figure 3.**
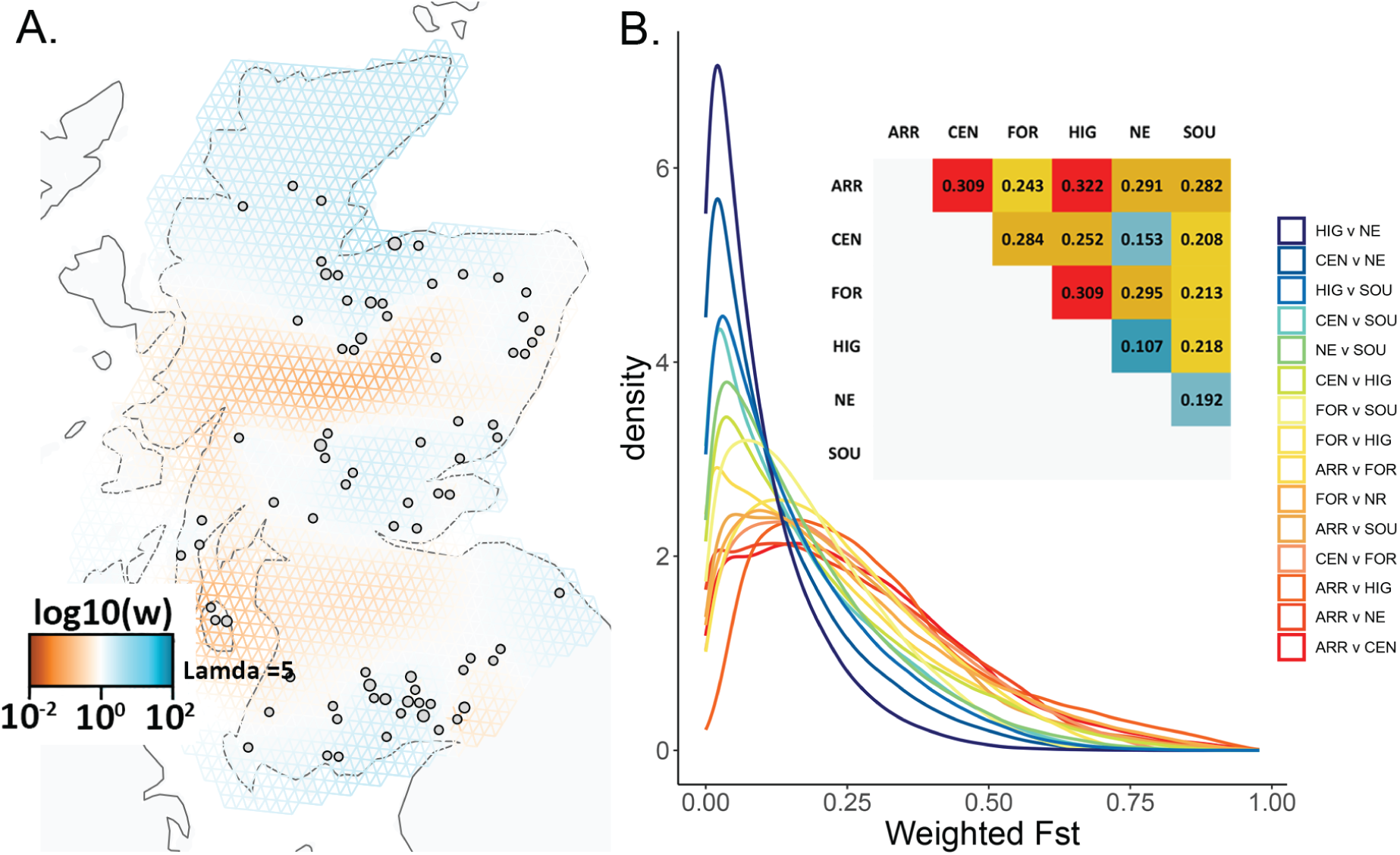
Migration and population differentiation (Fst). **A:** Fast estimation of effective migration surfaces (FEEMS) shows of low migration (brown) amongst corridors of higher migration (blue). Populations north and south of the Central Belt show no gene flow, while migration is also absent along the west coast and across tree-less areas. **B:** Weighted Fixation Index (Fst). Pairwise point estimates and genome-wide density plot of pairwise Fst. Population differentiation supports these migration patterns. Most populations show moderate to high differentiation, which is higher in the ARR and FOR pairwise comparisons. Log10(w) = relative effective migration

The fixation index (Fst), calculated as weighted pairwise point-estimates and genome-wide pairwise comparisons in 50kb windows (Fig.3), showed moderate to moderate-high differentiation between regions. Differentiation is higher in comparisons between the offshore Isle of Arran and isolated Formby populations and the mainland populations, being highest between Arran and the Highlands (Fst = 0.322) and lowest between the Highlands and the neighbouring north-east corridor (Fst = 0.107; Fig. 3).

Based on these findings, data were partitioned into six populations for further analysis – the Highlands (HIG), northeast (NE), central (CEN), southern (SOU), Arran (ARR) and Formby (FOR). The Highlands and NE were partitioned due to the unique admixture profile of the NE region, while Arran and Formby clearly do not constitute a single population. The SW and SE populations were combined into one southern (SOU) group as the most likely admixture scenario, but some geographical subdivision may be present. The three individuals from Argyll were removed from further population-level analyses due to ambiguous population status.

### 3.4. Genetic diversity and inbreeding

Genetic diversity, measured both by Watterson’s estimator (Fig. S7) and individual genome-wide heterozygosity (Figs. 4, 5), was exceptionally low in all individuals and across populations. Mean Watterson’s theta per population ranged from 2.57 x 10^-4^ (HIG) to 3.49 x 10^-4^ (SOU), although there were also numerous highly diverse genomic outlier regions associated with high-density SNP hotspots (Fig. S1), showing islands of high heterozygosity against a backdrop of low diversity. Mean genome-wide heterozygosity was some of the lowest reported for any species (Fig. 4), with a mean for the dataset of 2 x 10^-4^ and a range for individuals of 1.34 x 10^-4^ (individual from HIG) to 3.25 x 10^-4^ (individual from FOR; Fig. 5). The individual with higher coverage (22.5X) showed elevated heterozygosity compared to the same sample down-sampled to 5X. While this suggests an effect of depth on heterozygosity, the difference between these samples was 8.3 x 10^-5^, which is less than the intra-population range.

**Figure 4.**
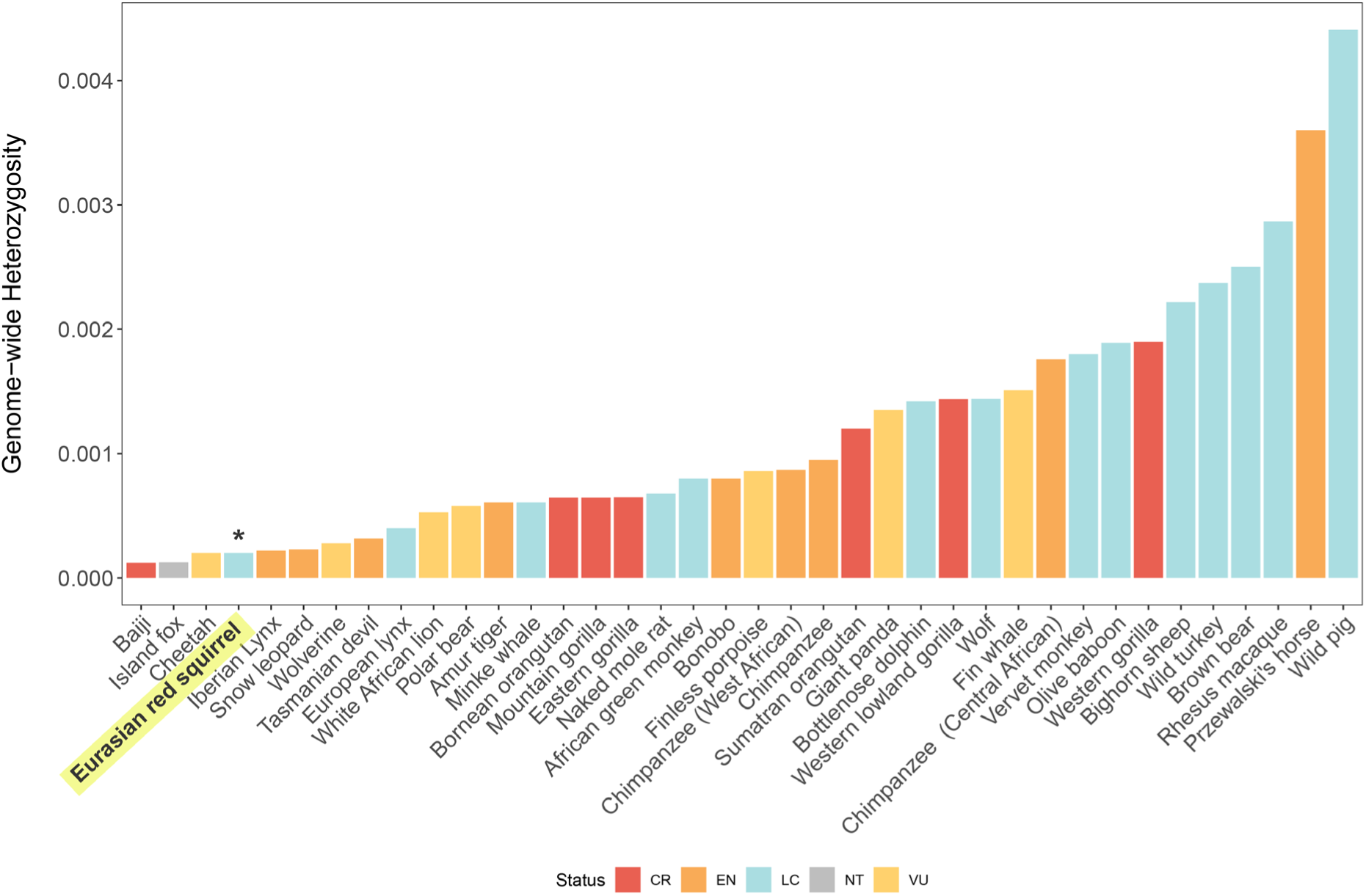
Comparison of genome-wide heterozygosity. Diversity measured by individual genome-wide heterozygosity for British (Scotland and Formby, England) red squirrels and other mammal species. IUCN Status (IUCN, 2023): CR = Critically Endangered, EN = Endangered, VU = Vulnerable, NT = Near Threatened, LC = Least Concern. Note that red squirrels are Endangered on the British Red List (Mathews et al., 2018). Data modified from (Robinson et al., 2016) and references therein.

**Figure 5.**
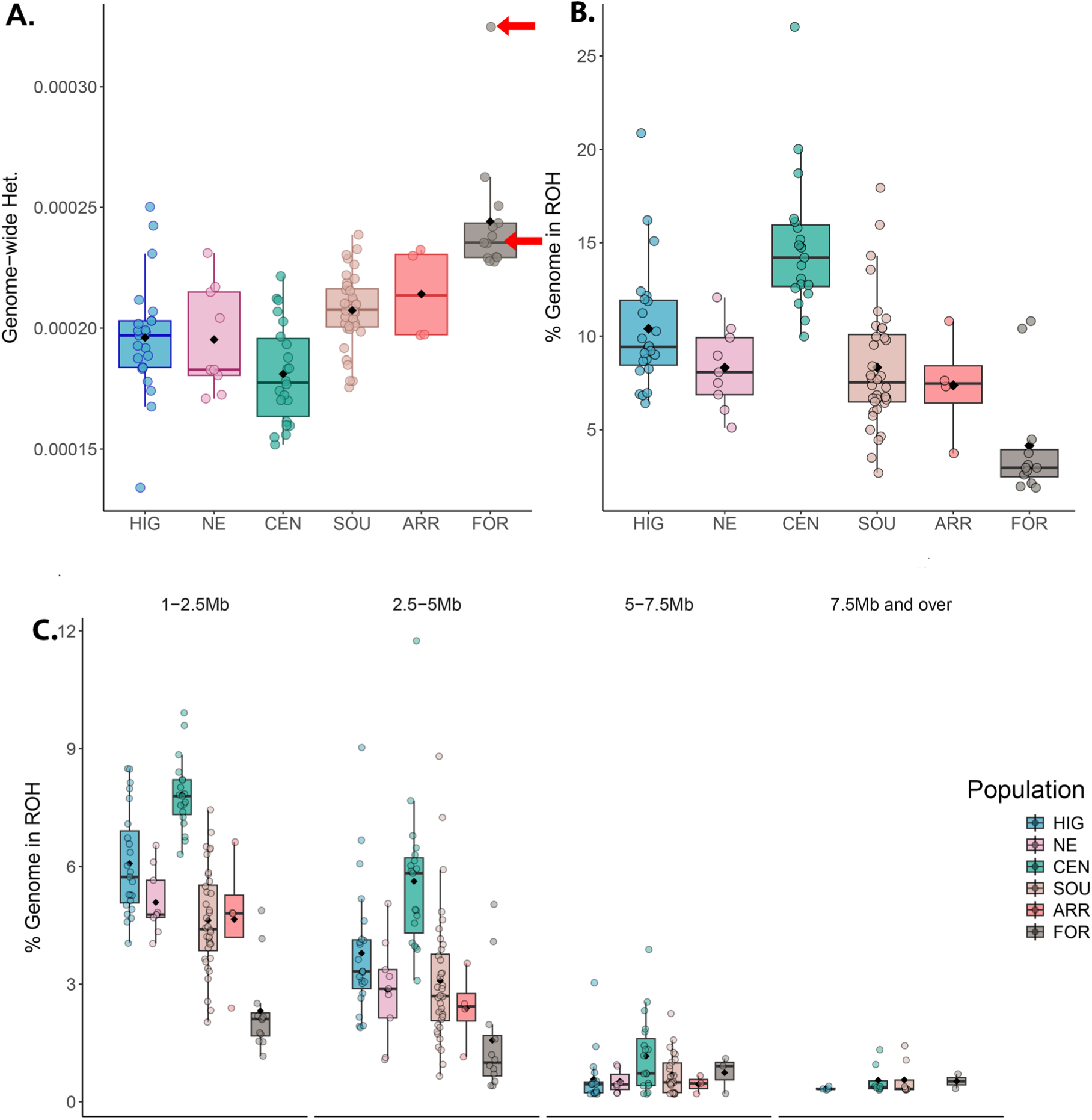
Individual genome-wide heterozygosity and runs of homozygosity. **A:** Boxplots of genome-wide heterozygosity for individuals and by population, where population means are indicated by a black bar. Heterozygosity is exceptionally and uniformly low across all populations, although highest in Formby. Individual heterozygosity was calculated twice for the high coverage individual from Formby at both high coverage (22.5X – top arrow) and downsampled (5X – bottom arrow). **B:** Percent of the autosomal genome in ROH and, **C:** Percent of the autosomal genome in ROH by size category, 1-2.5Mb, 2.5-5Mb, 5-7.5Mb, > 7.5M.

Comparison of genetic diversity inside and outside designated red squirrel strongholds was confounded by unequal sample sizes, and an uneven distribution of strongholds among squirrel populations (Fig. 1; Table 1, 2). Most strongholds are located in the Highlands and the west of Scotland, with few in the east, central and south-east regions (Fig. 1). For populations with adequate samples sizes (HIG, CEN and SOU), and for pooled data, two-tailed T-tests were performed to compare mean genome-wide heterozygosity. No significant differences in mean genome-wide heterozygosity were observed within and outside stronghold areas (Table 2), suggesting that heterozygosity is uniformly distributed across these areas.

**Table 2.** Two-tailed student’s T-Test, stronghold heterozygosity. T-tests, assuming unequal variances, were carried out within populations where there were sufficient sample sizes and for all data combined. No significant differences were observed.

|  | **n stronghold** | **n non-stronghold** | **t stats** | **t critical** | **p two-tail** |
| --- | --- | --- | --- | --- | --- |
| HIG | 18 | 5 | 1.743 | 2.45 | 0.132 |
| CEN | 4 | 15 | -2.234 | 2.306 | 0.056 |
| SOU | 8 | 28 | 2.402 | 2.776 | 0.074 |
| **All data** | **38** | **68** | **-1.235** | **0.221** | **0.968** |

The total number of ROH > 1Mb in the dataset was 11,547 and the average percentage of the genome covered by ROH (F_ROH_) was 9.48% (min 1.89%, max 26.57%; Table S3, Fig. 5). The CEN population showed highest mean % ROH genome coverage (14.87%), while the other populations had means between 5% and 10% and, notably, Formby had the lowest (4.15%). There was an abundance of short and moderate length ROH segments throughout the genomes (1-2.5Mb and 2.5-5Mb; Fig. 5), while longer ROH lengths (5-7.5Mb and > 7.5Mb) were rarer.

### 3.5. Tajima’s D and Demographic History

The Scottish mainland populations all showed moderately negative mean Tajima’s D (HIG = -0.878, NE = -0.447, CEN = -1.107, SOU = -0.980), while the isolated populations showed a negative, but less pronounced, departure from neutrality (ARR = -0.091, FOR = -0.098; Fig. 6). Negative Tajima’s D is generally indicative of an excess of rare variants associated with expansion and/or purifying selection, suggesting that some modest, recent, expansion may have occurred in the larger mainland populations, but not in the isolated populations of Arran and Formby. Interestingly, all populations showed large variations in Tajima’s D across genomic windows from strongly positive to strongly negative (Tajima’s D = ≥3≤-3), indicating differential effects of selection across the genome.

**Figure 6.**
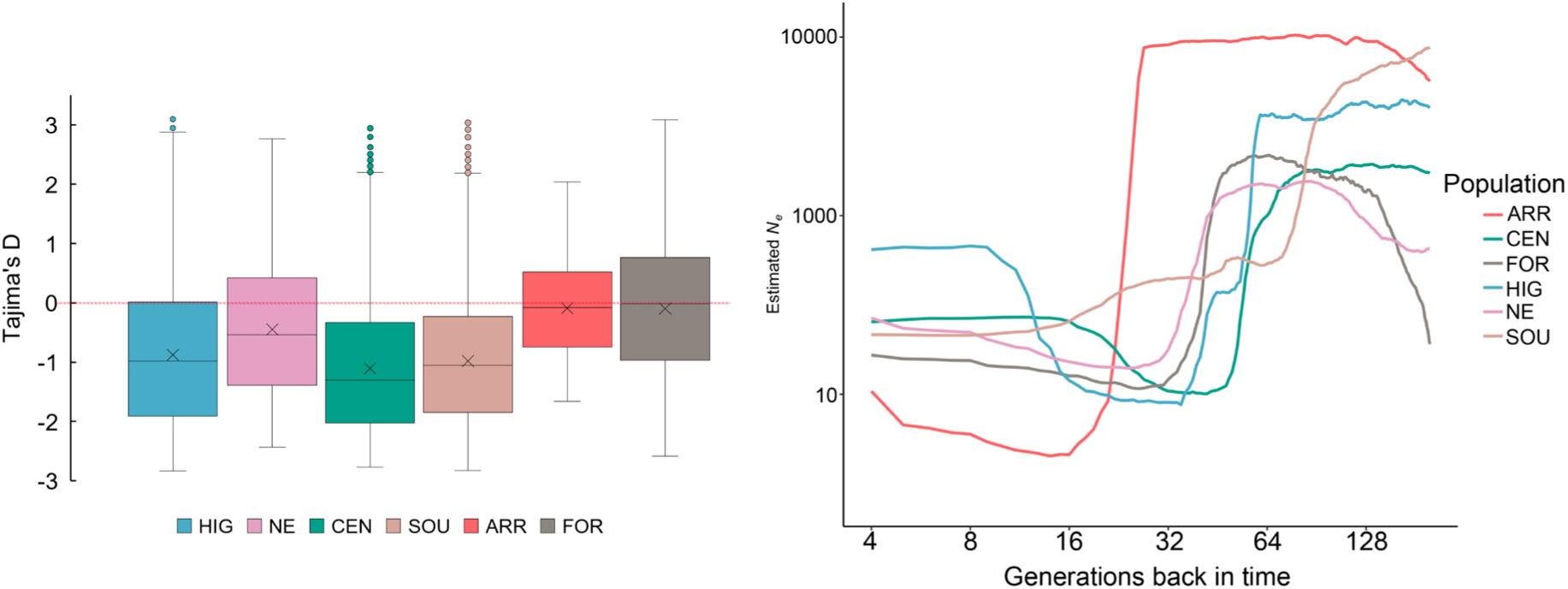
Genome-wide departures from neutrality and estimates of historical effective population size (Ne). **A:** Tajima’s D. Scottish mainland populations show a mean negative, moderate, departure from zero indicating possible recent expansion, while the isolated populations of Arran and Formby are close to zero. Tajima’s D was calculated in 50kb windows across the genome and outlying windows with strongly positive departures from zero can be seen in the larger populations of HIG, CEN and SOU. All populations showed a large range in positive to negative values across the genome. **B:** Historical Ne. Estimates were performed on 123,052 SNPs on a per-population basis. The analyses are accurate up to 200 generations, with Ne shown on a log scale. All population show a huge drop in Ne at different time scales, with the SOU population decrease beginning *c.* 100 generations ago, followed by FOR and mainland Scotland populations *c*. 64 gens and FOR c. 25 gens.

Estimates of historical effective population size (*Ne*) were performed over the last 200 generations using 123,052 SNPs (Fig. 6). There is a dramatic drop in *Ne* for all populations which is first observed in the South population *c.* 100 generations ago. This is subsequently observed in all mainland Scottish populations, and in the English Formby population, and is initiated *c*. 64 generations ago. The Isle of Arran population also experiences this phenomenon, although at a later date, with the commencement of the decrease *c*. 25 generations ago. All populations then spend *c*. 40 generations at very low *Ne*, with a small recovery observed *c*. 16 generations ago. Assuming a red squirrel generation time of three years, the declines for the mainland populations began *c.* 200 to 300 years ago (18^th^ to early 19^th^ centuries), with the Isle of Arran decline *c.* 75 years go (early to mid-20^th^ century).

## 4. Discussion

This study reports the first whole-genome analysis of the red squirrel, revealing the genetic legacy of past anthropogenic influences on remnant populations in a key region of conservation importance. Alarming and unexpected loss of diversity was observed across the genome and Scottish landscape, following an abrupt historic bottleneck. Previously undetected patterns of population structure and migration were highlighted, providing critical new information on spatial gene flow and habitat use. This is one of the most comprehensive genome-wide assessments performed to date of a population subjected to serial translocations and population fluctuations. The findings highlight the genetic risks associated with both anthropogenic threats and conservation efforts, which should be considered across species conservation management plans.

### 4.1. Genetic structure of red squirrel populations

Scottish red squirrel populations are composites of remnant indigenous Scottish populations with historical English and European additions. These populations have naturalised to the region over several centuries and now show geographically structured patterns of gene flow and population subdivision, which contrast with historic genetic lineages, presumably associated with founder source populations (Figs. 1, 2, 3). Contemporary structure is strongly influenced by natural and artificial landscape features, as well as founder composition and (presumably) the presence/absence of grey squirrel competitors (Figs. 2, 3). Spatial landscape modelling has previously suggested that the Cairngorms act as a barrier to squirrel movement (Slade et al., 2021), and this is clearly reflected in the genetic data (Fig. 3). The north-east (Aberdeen and Aberdeenshire) is a key contact zone for red squirrel populations and the east coast is the main corridor for latitudinal dispersal (Fig. 3). While urban development and grey squirrel presence has essentially isolated northern and southern Scottish red squirrel populations (Figs. 1, 3) this may, in fact, help to prevent intraspecies transmission of squirrelpox virus (SQPV), in which outbreaks have only been recorded in southern populations (McInnes et al., 2009); although the first case of squirrelpox north of the Central Belt has recently been identified (Wilson et al., 2024). The three individuals on the peninsula of Argyll and Bute that are genetically similar to the southern red squirrels (Figs. 1, 2) may have once been part of a population continuous with the southern population or, alternatively, could have been the result of unrecorded/unofficial translocations from the south into Argyll.

Red squirrel populations have disparate histories. Multiple historical introduction sites are recorded, but the origins of these animals are poorly documented, although England and Scandinavia have been identified as potential source areas (Harvie-Brown, 1881b, 1881a, 1880; Ritchie, 1920). The Formby population (NW England, Fig. 1) has been separated from other populations for many decades. It is reported to be founded by introductions from Europe in the early 20^th^ century (Gurnell and Pepper, 1993). The Arran population is undocumented in historical texts, and continental European haplotypes have been observed there (Ballingall et al., 2016; Barratt et al., 1999). This population is therefore also likely to be a 20^th^ century introduction from Europe rather than founded by natural colonisation. If Formby and Arran have genetic input from squirrels with common origins, this may explain their similarities in genetic admixture (Fig. 2). The inclusion of Formby as a comparative population highlighted a distinction between this population and the mainland Scotland groups. However, these results suggest that Formby may have a unique demographic history and not be representative of the now-extirpated English populations as a whole.

Notably, analyses of mitochondrial DNA did not detect any geographical structure (Fig. 2), as also found for British squirrels (Barratt et al., 1999). The mtDNA network may be reflecting historic lineage diversity that carries little phylogenetic signal in Scotland due to the disparate founder history. The mtDNA patterns observed here deviate from those typically observed in natural populations for other species, where networks tend to show haplotype frequencies that covary with geography (Dobigny et al., 2013; Vega et al., 2023). This lack of structuring is a strong indicator that human-mediated movement and translocations of individuals have occurred throughout the recent history of red squirrels in Scotland.

### 4.2. Genomic consequences of extreme founder effects

The timing and severity of the drop in effective population size (*Ne*, Fig. 6), is remarkably consistent with historical records (Harvie-Brown, 1881a, 1881b, 1880; Ritchie, 1920). The initial high *Ne* values are likely signals of ancestry from larger English and European populations, with the abrupt drop representing a severe bottleneck precipitated by founder effects (Fig. 6). The later founder effect, observed in the *Ne* plot for Arran (Fig. 6), reflects the later (20^th^ century) introduction date for the island compared to the mainland populations (18^th^ century). All populations, apart from the Isle of Arran, show a minimum estimated *Ne* of no less than 100 at their lowest point (Fig. 6). Some recovery has occurred, with most populations currently *c*. *Ne* = 100 – 500, and in the Highlands *c. Ne* = 900. The subsequent 20^th^ century introduction of grey squirrels has almost certainly suppressed any recovery process, and may be why *Ne* is now highest in the Highlands, where grey squirrels remain absent.

Red squirrel populations in Scotland exhibit some of the lowest heterozygosity reported for wild mammals (Fig. 4), and are comparable to species noted for extreme genetic impoverishment such as the Channel Island fox *Urocyon littoralis* (Adams and Edmands, 2023; Robinson et al., 2016), Iberian lynx *Lynx pardinus* (Abascal et al., 2016) and cheetah *Acinonyx jubatus* (Dobrynin et al., 2015) (Fig. 4). However, the red squirrel in Scotland does not have a comparably small population size; current estimates are *c.* 239,000 individuals (although this may be an over-estimate, Matthews et al., 2018) and historical records suggest significant expansion after restocking (Harvie-Brown, 1881a, 1881a, 1880; Ritchie, 1920). Some evidence of minimal post-bottleneck recovery is suggested by the negative Tajima’s D (Fig. 5) and slight improvement in *Ne* (Fig. 5). This clearly has not improved heterozygosity, which has persisted at very low levels over hundreds of generations despite historical population growth. This points to extreme founder effects with later expansions reversed and/or suppressed after the 20^th^ century introduction of the grey squirrel.

Interestingly, ROH did not indicate excessive recent inbreeding (Fig. 5). ROH represent stretches of the genome, where two alleles/haplotypes are identical by descent (IDB), indicating ancestry from the same ancestral haplotype and inferring levels of parental relatedness (Crow, 1954). Recombination and mutation break ROH so, in general, larger stretches are more characteristic of recent inbreeding than shorter lengths (Thompson, 2013). While there was large variability in F_ROH_ and the length of ROH (Fig. 5), inbreeding was generally characterised by short to moderate length ROH. This suggests that if there was an initial, post-bottleneck, accumulation of inbreeding tracts, they have subsequently been fragmented through local outbreeding, enabled by rapid demographic recovery typical of rodent species with short generation times.

### 4.3. Genomics informed management

Given that little increase in heterozygosity has been observed over the past three centuries, it seems unlikely that diversity will improve through natural population expansion. Long-term management interventions, such as conservation translocations among populations, or reinforcement from outside the region, may be required to improve the genetic status of red squirrels in Scotland. Enhancing physical population connectivity to increase gene flow is a recognised approach (Frankham et al., 2010; Gagnaire, 2020), and may prove beneficial to connect fragmented red squirrel populations. However, this should only be attempted in areas free of grey squirrels, as it risks facilitating the further dispersal of grey squirrels and, consequently, disease. Translocations of genetically selected, disease-free red squirrels between populations should be considered to increase diversity and promote gene flow. While these measures may come at the cost of dissolving population structure, they will promote a country-wide meta-population with improved genetic health.

Official strongholds are not evenly distributed throughout Scotland as a primary criterion for their placement is protection from grey squirrels (Slade et al., 2021). This species is present in south, central and east of Scotland; therefore, strongholds tend to be located in the north-west (Fig. 1). Genetic diversity was uniform within and out-with official stronghold areas (Table 2), but the distribution of strongholds means that they only fully capture genetic diversity from the Highlands (Table. 2, Fig. 1). Given the importance of the north-east and the east coast for red squirrel gene flow, creation of more strongholds in this area is desirable. Owing to the geography of the NE terrain, the north-east dispersal corridor is located at a physical bottleneck (Figs. 1 & 3), which has facilitated successful grey squirrel control in Aberdeen and Aberdeenshire (Tonkin et al., 2023). If grey squirrel suppression can be maintained, red squirrel strongholds in this region would help maintain one of the few natural red squirrel contact zones. From a genetic perspective increased stronghold presence in the central and south of Scotland would capture more country-wide variation. However, the viability of strongholds is unclear in areas densely populated with grey squirrels and with high SQPV prevalence. Considering the resource implications of maintaining multiple squirrel strongholds, further analysis of this dataset is warranted to gain a deeper understanding of the genetic diversity represented in each one and help inform future forest management. As an offshore island, the Arran stronghold (Fig. 1) has the advantage of being free from grey squirrels but at the cost of increased genetic isolation. Periodic additions of red squirrels from the disease-free mainland populations could ameliorate these issues.

## 5. Conclusions and wider implications

Red squirrels in Scotland exhibit extraordinarily low heterozygosity due to the genetic consequences of extreme historical founder effects, exacerbated by population subdivision and competition with an invasive species. Despite their poor genetic status, these represent the last substantial red squirrel populations remaining in Britain and should be regarded as a conservation priority. While the risk of outbreeding should always be considered when a planning conservation translocation (IUCN/SSC, 2013), in the case of the Scottish populations, where their history makes local genetic adaptation unlikely, the genetic benefits of translocating squirrels among populations, essentially creating a Scottish metapopulation, provide a strong argument for such an approach.

Interventions of this type are becoming ever more common as wild populations become increasingly fragmented and disjunct. Founder effects, exacerbated by population fragmentation, have been observed across species (Adams and Edmands, 2023; Colpitts et al., 2022; Kumar et al., 2023; Wilder et al., 2022), and many genetically depauperate species are now successfully managed using a metapopulation strategy. For example, Kenyan populations of the eastern black rhinoceros *Diceros bicornus micheali* are now wholly managed as a country-wide metapopulation (Amin et al., 2017), and notable success has also been recorded with the global management and subsequent re-introduction of the scimitar-horned oryx *Oryx dammah* (Humble et al., 2020; Ogden et al., 2020).

Future genomics-informed management of the red squirrel in Scotland would benefit from research that examines, *i*) evidence for harmful effects of low diversity, *ii*) post-translocation and post-bottleneck selection across the genome, *iii*) development of genetic tools for routine monitoring and *iv*) generation of, and comparative research with, genomic data from other UK and European populations. Until future management measures have impacted population genetic diversity, the use of any single Scottish population to provide all of the founders for introductions to other areas is strongly cautioned against. Many species contain a genetic legacy of past anthropogenic influence and the case of the red squirrel here illustrates that there is a need to take account of this, determine current surviving diversity and assess if intervention may be required.

## Supporting Information

**File 1**. svul_supp_mat.docx. Supplementary figures.

**File 2.** svul_SI_tabs.xlsx. Supplementary data and tables

## Acknowledgements

This research was funded by a Daphne Jackson Trust fellowship awarded to MMM and sponsored by the National Environment Research Council (NERC) and The University of Edinburgh. Additional contributions came from The University of Edinburgh Moray Endowment Fund, CryoArks, Forestry and Land Scotland and a Biotechnology & Biological Sciences Research Council (BBSRC) Institute Strategic Programme Grant BBS/E/D/30002276 to JS.

ACK acknowledges the support of the late Hon. Vincent Weir in supporting the preservation of the red squirrels in the NMS collection that were used in this study. ACK thanks the Negaunee Foundation for its support of a curatorial preparator who prepared some of the squirrels used in this study. We wish to extend our appreciation to the many members of the public, charities, volunteers and employed members that have submitted red squirrel carcasses to the R(D)SVS disease surveillance program over the last decade, without whom this research would not have been possible. The support and encouragement of the Daphne Jackson Trust to MMM is also gratefully acknowledged.

## Data Availability Statement

Genotype likelihoods, SNP genotypes and metadata can be assessed and downloaded from the Harvard Dataverse https://doi.org/10.7910/DVN/CK1ILL and will be made available 12 months from the date of publication. Code and scripts can be accessed at https://github.com/space-beaver/Red-squirrel-genomics-.git

## Ethics

This manuscript has not been submitted to another journal. All tissues used as part of this research originated from squirrels that died from natural causes or road traffic accidents; no live sampling was performed. Ethical approval for this project was gained on 05/08/2020 via the R(D)SVS veterinary ethical research committee (VERC) with approval reference 96.20.

## Conflict of Interest

The authors declare no conflicts of interest.

## Notes

### Competing Interest Statement

The authors have declared no competing interest.

