## Supplementary Figures for "Genomic insights into red squirrels in Scotland reveals loss of heterozygosity associated with extreme founder effects"

**Supplementary Material**

Melissa M. Marr^a^, Emily Humble^a^, Peter W. W. Lurz^a^, Liam Wilson^a^, Elspeth Milne^a^, Katie M. Beckmann^a^ , Jeff Schoenbeck^a^ , Uva-Yu-Yan Fung ^a^, Andrew C. Kitchener^b^, Kenny Kortland^c^, Colin Edwards^c^, Rob Ogden^a^.

^a^ Royal (Dick) School of Veterinary Studies (RDSVS) and the Roslin Institute, University of Edinburgh, Easter Bush Campus, Midlothian, Edinburgh, UK, EH25 9RG

^b^ School of Biological Sciences, The University of Hong Kong, Pokfulam, Hong Kong, 999077

^c^ Department of Natural Sciences, National Museums Scotland, Chambers Street, Edinburgh, UK, EH1 1JF and School of Geosciences, University of Edinburgh, Drummond Street, Edinburgh EH8 9XP, UK

^d^ Forestry and Land Scotland, Great Glen House, Leachkin Road, Inverness, UK, IV3 8NW

### List of Figures

- **Fig. S1.** SNP densities across the red squirrel genome.
- **Fig. S2**. Folded site frequency spectra (SFS).
- **Fig. S3**. PCA variance in the first 15 PC’s
- **Fig. S4.** PCA plot for pruned sample dataset.
- **Fig. S5.** NGSadmix likelihood curve and DeltaK for the admixture analysis.
- **Fig. S6.** Fast estimation of effective migration surfaces (FEEMS).
- **Fig. S7.** Watterson’s estimator

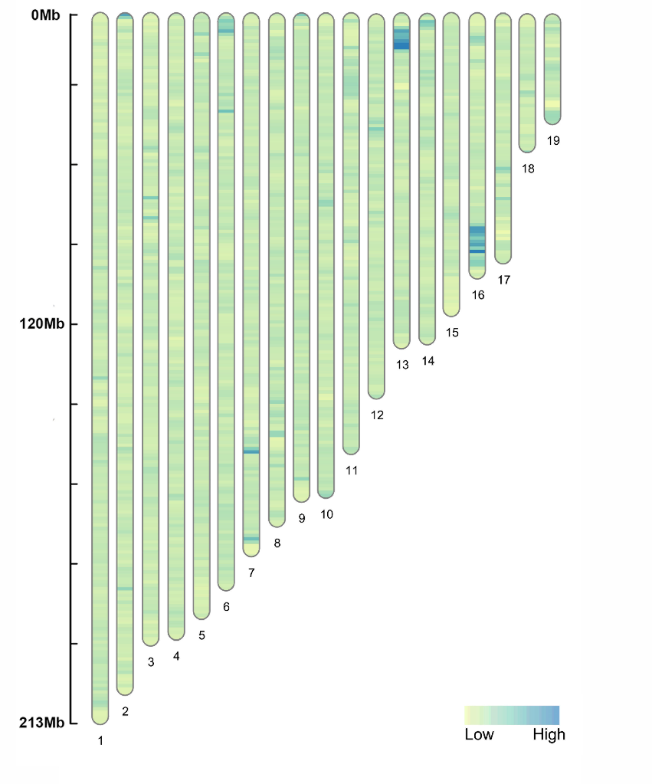

**Fig S1. SNP densities across the red squirrel genome**. Counts are binned in 1Mb windows across the 19 autosomes of the *Sciurus vulgaris* reference genome (GCA_902686455.2 (Mead et al., 2020)) for the low coverage ANGSD run. Small SNP density hotspots can be seen throughout the genome, with two large hot spots on chr13 and chr16.

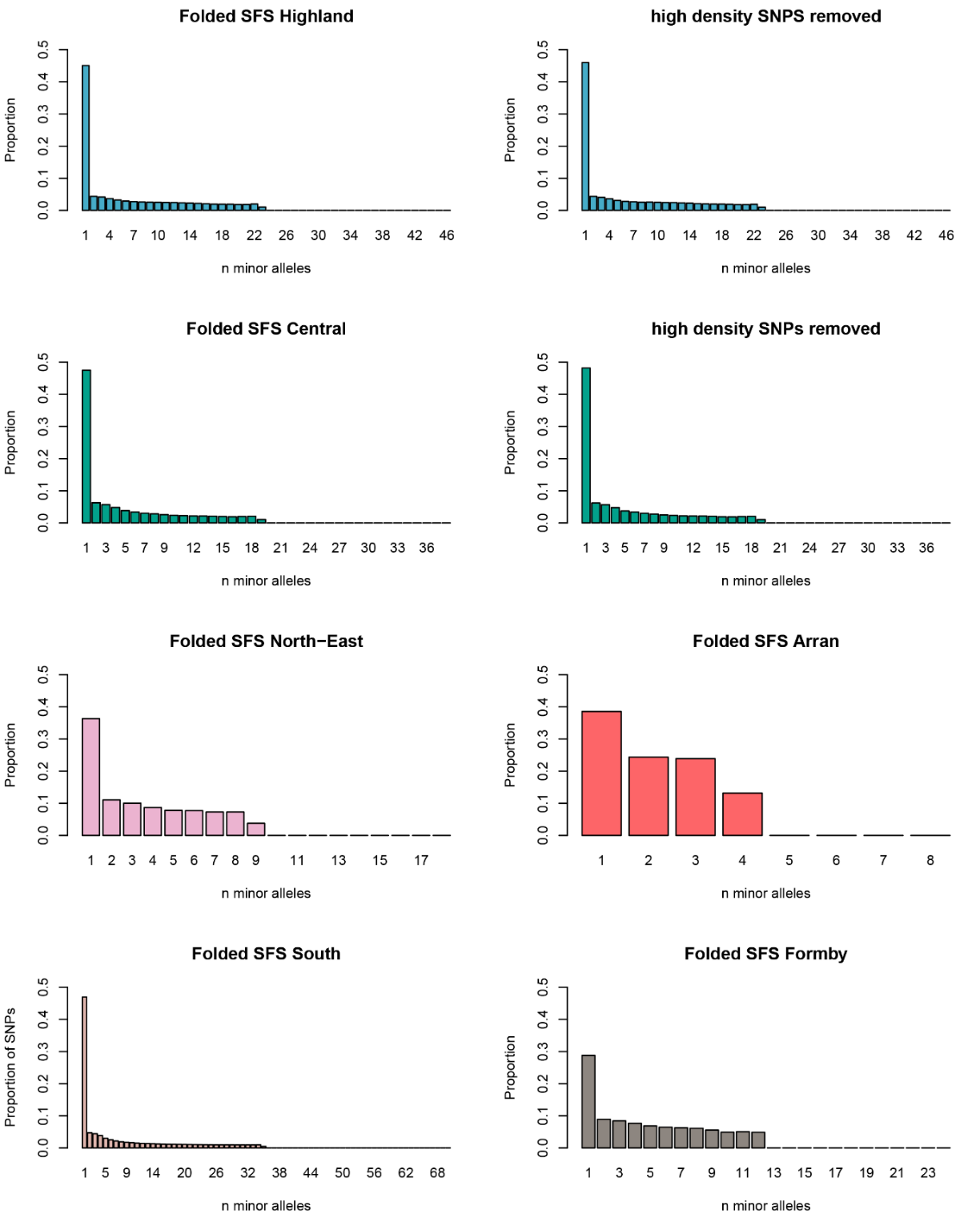

**Fig S2. Folded site frequency spectra (SFS).** Proportion of minor alleles per frequency class for each population. Histograms indicate an excess of rare variation in most populations. The absence of high numbers of rare alleles in the smaller, isolated populations of Arran and Formby may indicate population stasis or be an artifact of small sample size. Folded SFS for the Highland and Central populations are also shown without the high-density SNP hotspots areas of chrs 13 and 16; no discernible effect can be seen.

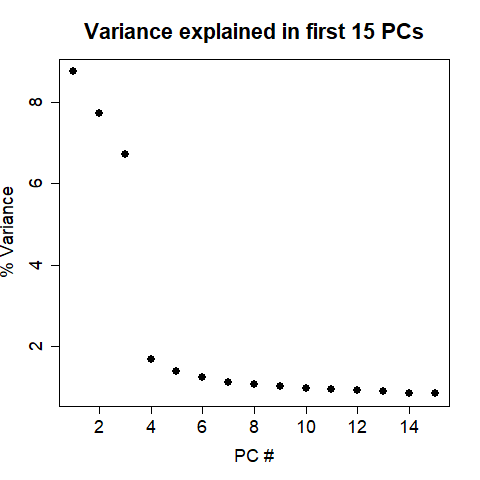

**Fig. S3. PCA variance in the first 15 PC’s.** Variance explained by the first 15 Principal Components (PCs) for the full sample dataset *n* = 106.

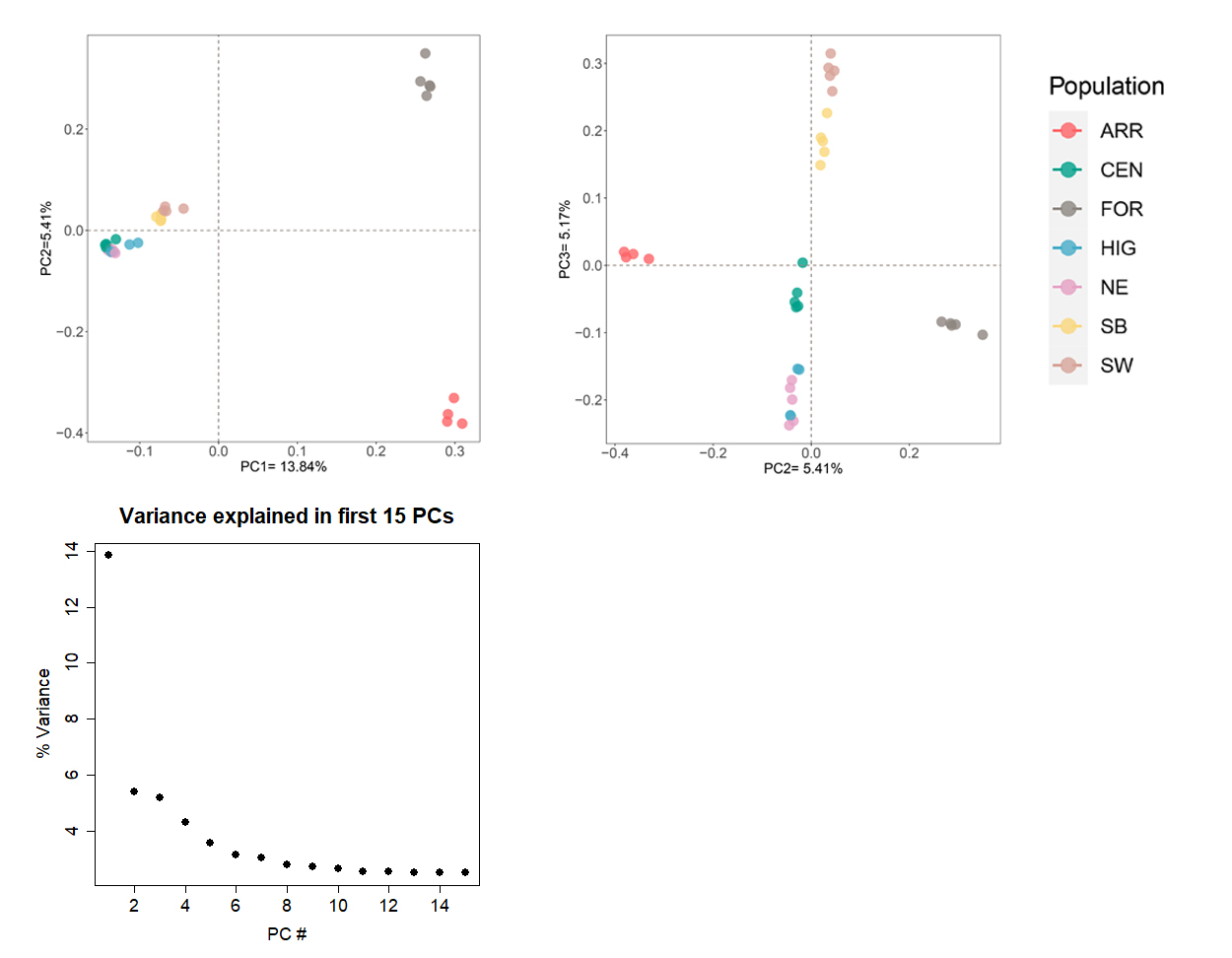

**Fig. S4. PCA plots, and PC variance, for pruned sample dataset**. PC’s 1v2 and 2v3 are shown with sample size *n*=4 or *n*=5 per population. Random subsampling for equal sample sizes per population showed the same subdivision into islands v. mainland across PC1 and mainland divisions between the Highland + NE, Central and South groups. The Isle of Arran and Formby did not group together across PC2 in these plots.

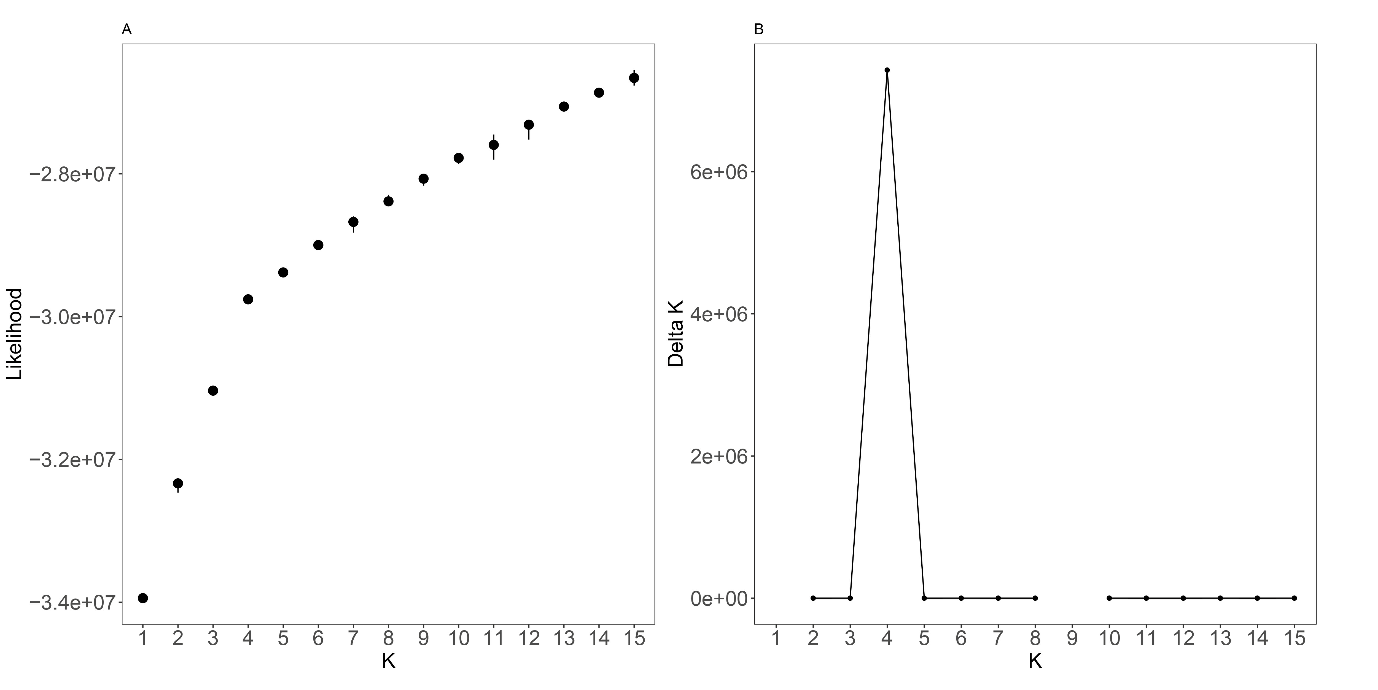

**Fig. S5. NGSadmix likelihood curve (A) and DeltaK (B) for the admixture analysis.** The analysis was run for K=2-20. Both the likelihood plot and DeltaK strongly indicate K=4 as the most likely number of populations.

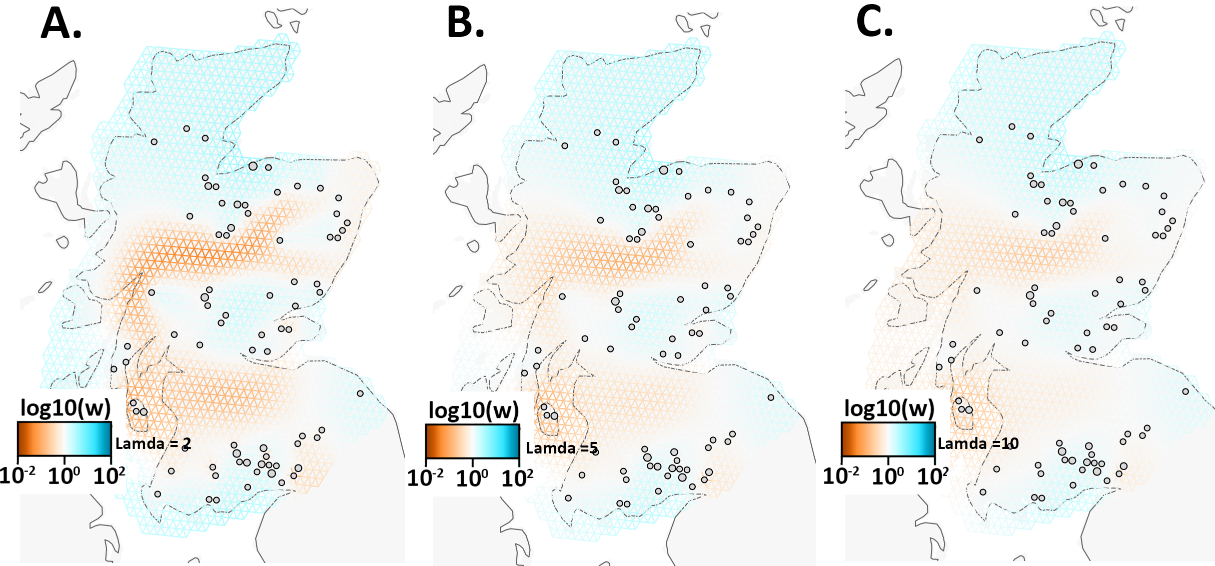

**Fig. S6. Fast estimation of effective migration surfaces (FEEMS).** Plots are shown for λ = 2, 5 and 10, showing increased structure with reduced value for lambda. Brown indicates higher migration rates. Log10(w) = relative effective migration.

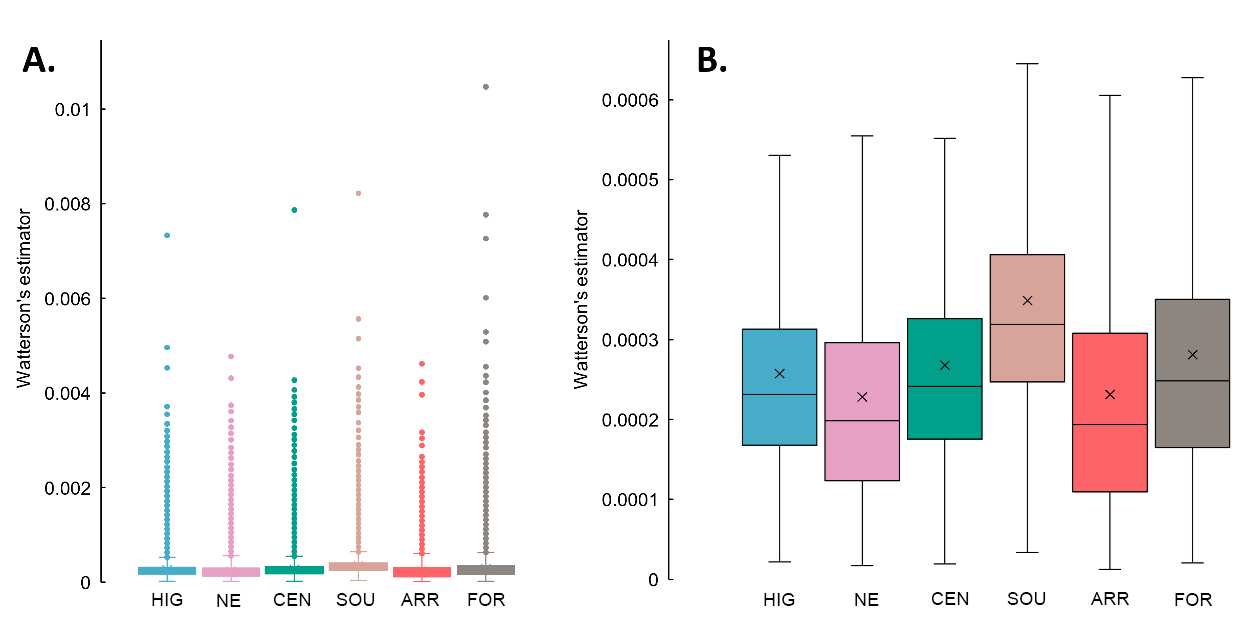

**Fig. S7.** **Watterson’s estimator (theta).** Theta was calculated in 50kb windows with a 10kb step size. Mean theta was extremely low for every population. Hot spots of heterozygosity can be observed across the genome (A), many of which are windows across high density areas on chr13 and chr16 and other SNP dense regions. Removal of these outlier regions (B) shows higher resolution of population means and interquartile range.
